# Deep learning models for lipid-nanoparticle-based drug delivery

**DOI:** 10.1101/2020.04.06.027672

**Authors:** Philip J Harrison, Håkan Wieslander, Alan Sabirsh, Johan Karlsson, Victor Malmsjö, Andreas Hellander, Carolina Wählby, Ola Spjuth

## Abstract

Large-scale time-lapse microscopy experiments are useful to understand delivery and expression in RNA-based therapeutics. The resulting data has high dimensionality and high (but sparse) information content, making it challenging and costly to store and process. Early prediction of experimental outcome enables intelligent data management and decision making. We start from time-lapse data of HepG2 cells exposed to lipid-nanoparticles loaded with mRNA for expression of green fluorescent protein (GFP). We hypothesize that it is possible to predict if a cell will express GFP or not based on cell morphology at time-points prior to GFP expression. Here we present results on per-cell classification (GFP expression/no GFP expression) and regression (level of GFP expression) using three different approaches. In the first approach we use a convolutional neural network extracting per-cell features at each time point. We then utilize the same features combined with: a long-short-term memory (LSTM) network encoding temporal dynamics (approach 2); and time-series feature extraction using the python package tsfresh followed by principal component analysis and gradient boosting machines (approach 3), to reach a final classification or regression result. Application of the three approaches to a previously unanalyzed test set of cells showed good predictive performance of all three approaches but that accounting for the temporal dynamics via LSTMs or tsfresh led to significantly improved performance. The predictions made by the LSTM and tsfresh applications were not significantly different. The results highlight the benefit of accounting for temporal dynamics when studying drug delivery using high content imaging.

## Introduction

Automated microscopy imaging, enabling multiple parameter measurements from a broad range of samples, is one of the most powerful tools for investigating complex biological processes, such as drug delivery, disease biology, and potentially toxic off-target effects. Current image-based assays however tend to either focus on a single time point or interrogate biological systems over time using relatively simplistic methods (such as comparing the first and last time points). The importance of looking at cellular systems as ongoing processes is apparent and such experiments almost always lead to new biological or pathological insights (1).

In this paper we explore temporal cellular responses to RNA-based therapeutics. Although such therapies have shown significant promise, more research into the RNA delivery will be necessary to realise their full potential. One of the most clinically relevant methods for RNA delivery utilizes lipid-nanoparticles (LNPs) (2), which can be tracked over time using automated microscopy systems. Time-lapse microscopic imaging data has a number of challenging attributes including high information content (although this is often sparsely distributed across the images), high dimensionality and temporal components. To address these issues, the current work uses two strategies. First, the effect of a nanomedicine on any given cell becomes evident only at later time points, when protein production finally reveals previous effective mRNA delivery by the nanoparticles. At this point it is however too late to evaluate how this delivery occurs. It is thus necessary to follow each cell over time. We hypothesize that it is possible to predict if a cell will produce GFP based on cell morphology at time-points prior to GFP expression. If cells can be accurately classified at early time points, using their appearance, then mechanistic information may be extracted. For example, the configuration and content of the endosomal network in cells that will produce GFP can be compared to cells that will not, revealing how cells are processing nanoparticles that successfully delivered mRNA. This information can be used to engineer particles that favour this process, resulting in more effective therapies. A second strategy is to optimise this process to work with relatively small datasets. Small data sets have many advantages (faster acquisition times, reduced storage requirements, decreased processing time) but are more challenging when used with deep learning approaches that typically require much larger datasets.

In biomedical image analysis the application of convolutional neural networks (CNNs) has yielded significant improvements in the automation of classification, segmentation, feature extraction and restoration of image data (3). A major bottleneck when applying these deep learning methods to cell and tissue images is the scarcity of annotated data. Studies have shown that reusing models trained for other tasks alleviates this problem (4). This is known as transfer learning - the transfer of knowledge between tasks - and is often beneficial when a limited amount of annotated data is available, such as in image cytometry, where manual annotations are time-consuming to acquire and require a high level of expertise to make. Furthermore, deep learning models trained on biomedical images, captured under specific experimental conditions and imaging setups, can have poor generalizability. To overcome these limitations, large annotated datasets, like ImageNet (5), can be used to pretrain state-of-the-art deep learning models. The transferred parameter values, providing good initial values for gradient descent (the most common optimization algorithm for learning the weights/parameters in neural networks), can be fine-tuned to fit the target data (4).

The sequential information contained in time-lapse imaging (i.e. information that is not available from a single time point image) can be of great utility for shedding light upon complex temporal cell processes, such as division, differentiation, and migration (6). Recurrent neural networks (RNNs) are a popular choice for analyzing sequential data and are used ubiquitously in machine translation and speech recognition (7) and in the field of biology they have been used for functional quantification of DNA sequences (8) and for the subcellular localization of proteins (9). The basic RNN consists of hidden states that evolve over time though nonlinear functions of the previous hidden states and the current inputs. When viewed across time RNNs are akin to very deep feedforward networks; as such they can be difficult to train (suffering from vanishing and exploding gradients) and can fail to retain information from the more distant past (10). Various solutions to these problems have been proposed, such as skip connections across time. The most successful solution is the long-short-term memory (LSTM) network (11), which controls the flow of information through gate units, such as the “forget gate”, an adaptive version of which enables the LSTM cell to reset memory blocks whenever their content becomes outdated (12).

RNNs automatically extract time-series features from temporal data relevant for the classification or regression task at hand. Alternatively one can extract such features directly from the time-series prior to performing classification or prediction. There exists a number of such (traditional) features developed across a wide array of scientific disciplines (13); such as measures based on fluctuations (physics), autoregressive methods (economics), and entropy (medicine). The python package tsfresh (14) contains a comprehensive set of these features useful for machine learning models, computing a total of 794 time-series features (based upon 63 time-series characterization methods).

The early classification of time series, whilst maintaining suitable levels of accuracy in the predictions, is of significant interest in many scientific domains (15). Furthermore, if we can predict the final outcome from the changes in the cells at early time points we can also explore the biological mechanisms responsible for successful uptake. However, before exploring these underlying biological mechanisms we first need to know if we can make such predictions at the cell-level and if so which approach is most suitable for modeling the cell dynamics. This is the focus of the current manuscript.

## Materials and Methods

### Nanoparticle production

The lipid components of the nanoparticles were obtained from various sources: the ionizable cationic lipid O-(Z,Z,Z,Z-heptatriaconta-6,9,26,29-tetraem-19-yl)-4-(N,N-dimethylamino)butanoate (DLin-MC3-DMA) was synthesized at AstraZeneca, the 1.2-distearoyl-sn-glycero-3-phosphocholine (DSPC) and 1.2-dioleoyl-sn-glycero-3-phosphoethanolamine-N-(lissamine rhodamine B sulfonyl) was obtained from Avanti Polar Lipids, 1.2-dimyristoyl-sn-glycero-3-phosphoethanolamine-N-[methoxy(polyethyleneglycol)-2000] (DMPE-PEG2000) was obtained from NOF Corporation, and cholesterol was obtained from Sigma–Aldrich. The nanoparticle cargo, modified (“CleanCap”), cyanine-5 labelled mRNA coding for enhanced green fluorescent protein (GFP), was obtained from Trilink. Citrate buffer was purchased from Teknova, and HyClone RNase free water was obtained from GE Healthcare Cell Culture.

Lipid components were dissolved in ethanol, and GFP mRNA was dissolved in an aqueous solution (pH 3.0 citrate buffer). These solutions were rapidly combined, at a 3:1 ratio, using a NanoAssemblr microfluidic mixer, from Precision NanoSystems Inc., resulting in a 3:1 molar ratio (10:1 weight ratio) between the cationic lipids and the mRNA nucleotides. The final particle composition was 50% Dlin-MC3-DMA, 9.9% DSPC, 38.5% cholesterol, 1.5% DMPE-PEG and 0.1% DOPE-rhodamine (molar percentages). The resulting LNPs were dialyzed overnight to normalise the pH and remove ethanol, using Slide-A-Lyzer G2 dialysis cassettes from Thermo Scientific with a molecular weight cutoff of 10 K. The resulting lipid particles have an orange-fluorescent lipid body, and a red-fluorescent mRNA cargo that codes for GFP.

Following micromixing and dialysis the size of LNPs was evaluated using dynamic light scattering ( Zetasizer Nano ZS from Malvern Instruments Ltd.). The encapsulation and concentration of mRNA were determined using a RiboGreen assay (mRNA inside the particles is not accessible to the dye). The encapsulation in all of the samples was typically 90–99%. Lipid-nanoparticles (LNPs) were manufactured less than 1 week before dosing.

### Cell culture

One day prior to imaging, HepG2 cells (a hepatocellular cell line) were seeded (in DMEM low glucose without phenol red (ThermoFisher) supplemented with 1% human serum, non-essential amino acids and 10 mM HEPES (all from Gibco)) into 384-well CellCarrier Ultra plates (Perkin Elmer) that had been pre-coated with rat collagen (Gibco). The cells were pre-stained using a 30 min. incubation with CellTracker Deep Red (ThermoFisher) and washed before fresh medium (as above) was added. Immediately before the start of the time-lapse imaging LNPs were added to triplicate wells in an equivalent volume of a 2x particle solution made using the same cell culture medium.

### Automated time-lapse imaging

All wells were imaged using a Yokogawa CV7000 robotic confocal microscope that maintained cell culture growth conditions (37 °C, 5% CO2, 100% humidity). Two fields of view, at every time point, were imaged using a 40x objective (0.95 air, Olympus), and two different imaging modes (brightfield and confocal fluorescence). Fluorescence images were in turn acquired (at all time points and fields of view) using minimal exposure times and three different spectral combinations (emission/excitation nm light) for green (488/525), orange (561/600) and red (640/676) fluorescence. The red fluorescent cellular counterstain does overlap the red fluorescence of the mRNA cargo but these signals can be separated morphologically because the mRNA fluorescence is granular or punctate.

For each well two fields of view were imaged (each 2554 x 2154 pixels with a pixel resolution of 0.1625 *μ*m/pixel). Images were acquired every ten minutes for twelve hours (72 time points). In Figure 1 we show example images from our time lapse experiment from one field of view in one well. Our experiments were done in triplicate with three LNP doses: 0, 31.6 and 316 ng mRNA/well (25ul). In Figure 2 we show the number of LNP positive spots per cell area over time and the mean GFP intensity over time. For our cell-level models (detailed below) we used as input three of the imaging channels - CellTracker counterstain (CS), brightfield (BF) and LNP channels - to predict the final GFP result, derived from the GFP channel at *t* = 72. As GFP expression does not begin until around the twentieth time point, but the LNP uptake occurs rather rapidly prior to this time point (see Figure 2), we were interested to see if we could make GFP predictions based on these three input channels during the early time points (*t* = 1 to *t* = 20).

**Figure 1:**
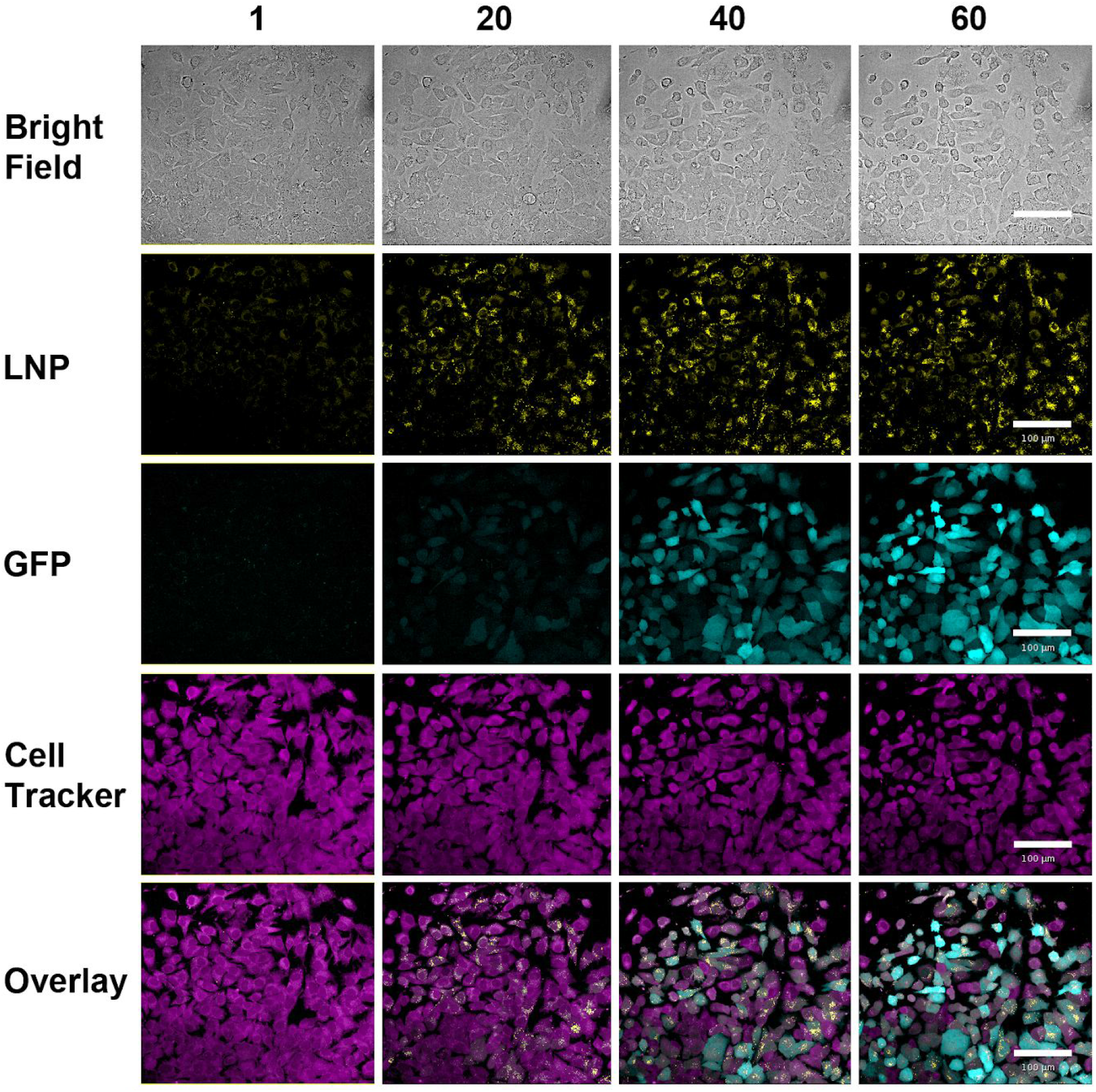
Representative images from cellular time-lapse imaging series showing time points 1 (~0h), 20 (3.75h), 40 (7h) and 60 (10.25h), of 72 time points, in columns. Rows show brightfield and false color confocal images of cellular LNP and GFP accumulation over time as well as the cell tracker counterstain. All images are shown with identical contrast settings. Scale bars show 100 *μ*m.

**Figure 2:**
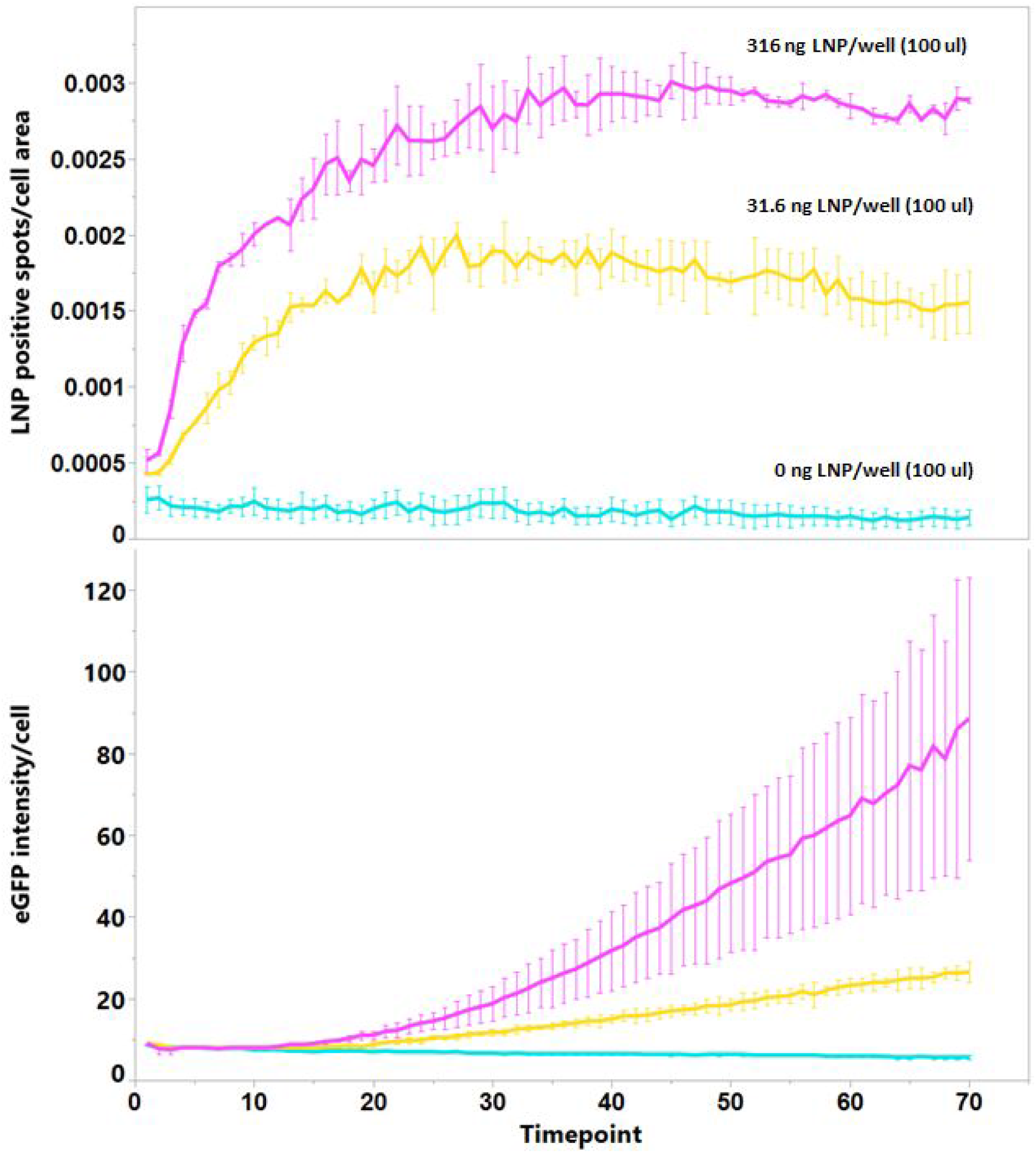
(a) mean number of LNP positive spots/cell area ∓ sem and (b) mean eGFP intensity/cell ∓ sem for the three mRNA doses (0, 31.6 and 316 ng mRNA/well where wells contain 100*μ*l cell culture medium with 1% fetal bovine serum). One time point is 10 minutes so time point 20 is about 3.5 hours.

### Cell segmentation and tracking

The cell centres were identified manually in the first time frame, and around each center point a 200 x 200 pixel window was drawn (the window dimensions being set based on the size of the cells). To delineate the cell nuclei, watershed segmentation (WS) (16) was used on the CS imaging channel. The WS segmentation algorithm views the image from a topographical standpoint (a 3D representation with intensity on the z-axis) and the pixels as belonging to one of three categories: (i) minima, into which water would drain; (ii) those of a catchment basin (where water would drop to a single minima); and (iii) those where water could go to more than one minima. This third category of pixel form connected paths, the watershed lines, which provide the desired segmentation. A significant benefit of this method is that it can incorporate knowledge based constraints, in the form of markers, to aid the segmentation. Internal markers, or seed points, define the minima in a given region and external markers define the areas into which the segmentation cannot pass (16). As our cells do not move considerably during the ten minutes between each time point we used the center point from the segmentation at time *t* as the internal marker for the WS algorithm at time point *t* + 1. The 200 x 200 pixel window edges were used as external markers.

### Cell-level time-lapse data

The primary purpose of the segmentations detailed above was to enable accurate tracking of the individual cells and to provide the cropping windows for our subsequent deep learning models, giving a dataset containing 72 time points with 192×192 pixel windows cropped around each cell (this value of 192, slightly smaller than the segmentation windows above, being selected to give an even flow of the images through the convolution and pooling operations of the neural networks). Any cell whose centre point became less than 96 pixels from the edge of the microscope’s field-of-view during the duration of the experiment was discarded. One of the high LNP dose wells had three time points of missing data early in the experiment. This caused difficulties for our tracking algorithm and missing data for our time-series models and hence this data was also excluded. After these removals we were left with time-lapse data for 774 cells.

The cells were randomly shuffled and approximately 80% were used for model training (and validation) and the rest for testing. We used as the target in our models the sum of the pixels in the GFP channel in the center of the 192 x 192 crop (a smaller window of 96 x 96 pixels) - this was done to avoid contamination by other potentially GFP-expressing cells close to the focal cell. We performed the modeling in both classification and regression mode. The threshold for dividing the positive and negative cells in classification mode was determined based on the maximum GFP pixel sum for the cells in the control well (i.e. those in the wells with no mRNA administered). This procedure gave 602 cells in the training set (387 positives and 215 negatives) and 172 cells in the test set (123 positives and 49 negatives).

### Modeling approaches

For classification, we predict successful/unsuccessful GFP expression, and for regression the actual amount of GFP expressed is predicted. The threshold for successful GFP expression was set to be greater than the maximum sum across the center crop of GFP windows for the control cells. The GFP value in the regression-based models was transformed on the natural logarithm scale and centred on zero (this standardization method giving better results than alternative methods, based on preliminary explorations with one of the training cross-validation folds). We used 5-fold cross validation on the training set. One issue with deep learning, and machine learning in general, is the problem of overfitting, whereby the neural network can learn an imprint the entire dataset (3), essentially memorising the data, such that rather than learning generalizable features, the network simply learns the features plus the noise in the training data. To overcome this problem a validation set (a subset of the training data) is required with which to evaluate/monitor the trained model. Such validation is essential for selecting the appropriate network architecture and its hyper parameters (such as the number of neurons in a given layer (3)). A final evaluation of the model on unseen data, a test set, is then required - data which has not been seen before, neither during (proper) training nor validation. We also used data augmentation (for both the training and test sets), via mirroring and 90 degree rotations, to give 8 times as much data. For the time series modeling the same augmentation was used across the time points for each time-series data points.

Schematic diagrams of the modeling approaches detailed below are given in Figures 3 and 4. In Figure 3 the framework that uses a combination of CNNs (for feature extraction) and an LSTM for time-series modeling is shown. In Figure 4 an equivalent framework is shown where the time series modeling is performed via time series feature extraction (tsfresh, detailed later) followed by principal components analysis (PCA) for dimension reduction and gradient boosting machines (GBMs; (16)) for the final classification or regression result.

**Figure 3:**
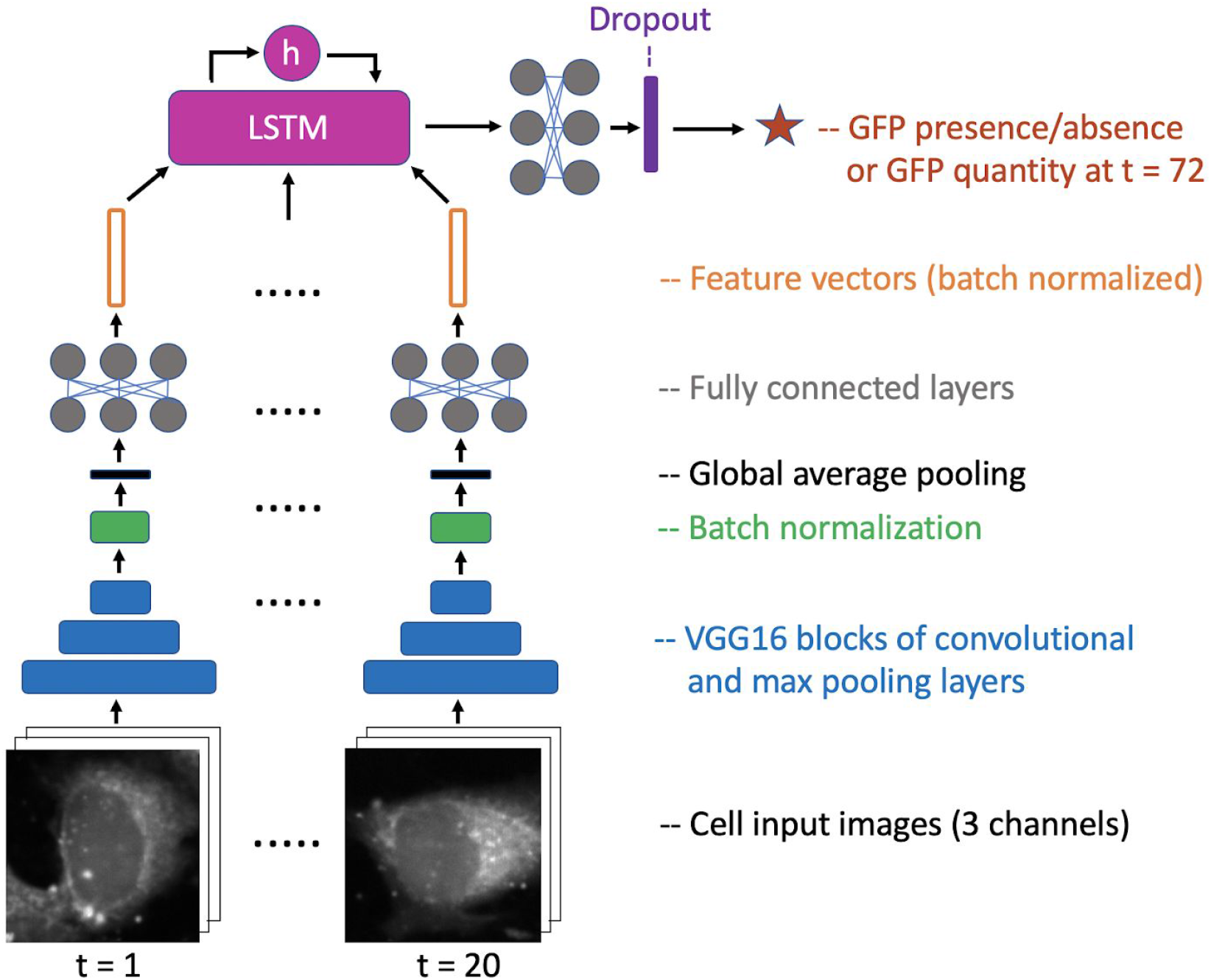
Modeling schematic combining the CNN (for feature extraction) with the LSTM. Figure inspiration from (24). The input images being the three imaging channels (CS, BF and LNP) across the first twenty time points of the experiment and the final target being the presence or absence of GFP in the final time point (t = 72) when in classification mode and the actual amount of expressed GFP for the cell when in regression mode. The feature vectors were of size 32.

**Figure 4:**
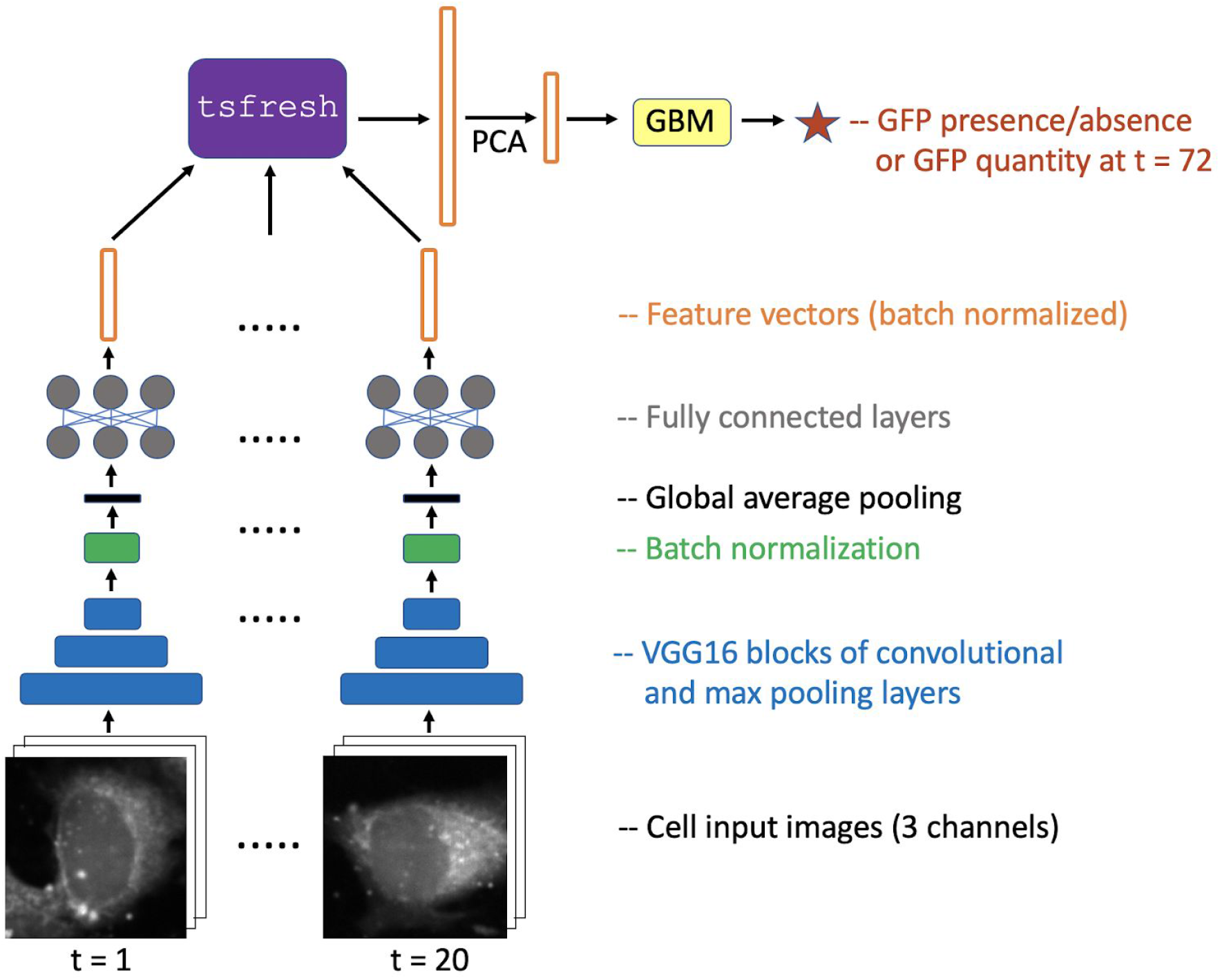
Modeling schematic combining the CNN (for feature extraction) with tsfresh (for time series feature extraction) followed by PCA and gradient boosting machines (GBMs). The input images being the three imaging channels (CS, BF and LNP) across the first twenty time points of the experiment and the final target being the presence or absence of GFP in the final time point (t = 72) when in classification mode and the actual amount of expressed GFP for the cell when in regression mode. The feature vectors were of size 32.

### Convolutional Neural Networks

We used CNNs to extract features from the cells and time-series modeling to take account of the evolution of these features through time. The CNNs were trained on all the cell data across the first 20 time points. This was done in part to provide more training data for the CNNs and also to ensure that the features extracted from the penultimate dense layer of the CNN (of size 32) would correspond to the same features for each time point (a necessity for the time-series models). Hence, the CNNs were trained using all the data across the first twenty time points, but without knowing explicitly from which time point a particular image came.

As our dataset is relatively small we used transfer learning via a base model in the CNNs, specifically we utilized the convolutional and pooling layers from the first three blocks of the VGG16 model (17) trained on ImageNet, except for the final pooling layer of the third block. Atop this base model we added the following layers: batch normalization; global average pooling; a batch normalized dense layer of size 32 (this being our latent feature vector for the time series models); followed by a RELU activation and a final dense layer of size 1 (with a sigmoid activation when in classification mode and no activation when in regression mode). Hence, prior to the final dense layer, the CNN has the same structure as we have before the LSTM and tsfresh parts shown in Figures 3 and 4. Since our classes were somewhat imbalanced (for classification) weighted cross entropy was used. For regression we utilized mean squared error (MSE) loss. We used a batch size of 32 and the Adam optimizer. Using one of the five cross validation folds we performed explorations to determine an upper limit on the number of epochs to train for and the learning rate(s) to use (0.001, 0.0001 or 0.00001). The models were first trained with the base frozen and then with the base unfrozen. These settings were then used across all five folds and subsequently the weights from the best epoch (in terms of validation loss) for each fold were saved.

### Time-series modeling

We explore two alternative methods for the time series modeling: either recurrent neural networks (LSTMs); or time-series feature extraction (tsfresh) followed by PCA (for dimension reduction) and GBMs, for the final prediction.

#### LSTMs

For our RNN based implementations (see Figure 3) we used LSTMs. To determine the appropriate architecture and hyper parameters for the LSTMs we drew 200 samples from a randomized grid of possibilities for each of the five folds. The following values defined the grid: whether or not to use a bidirectional LSTM; the number of LSTM layers (1 or 2) for the non-bidirectional LSTMs; whether or not to include a penultimate dense layer (after the LSTM and prior to the prediction); the number of neurons for each layer (32, 64, 128 or 256) and; the dropout rates for the LSTM layers and the penultimate dense layer (0.1, 0.2 or 0.5). In all cases a batch size of 32 was used. Again the learning rates and maximal number of epochs to train for were selected by explorations using one of the cross-validation folds.

#### tsfresh

As an alternative to using RNNs we explored extracting time-series features from the 32 dimensional feature vectors across the twenty time points using tsfresh. tsfresh first extracts the features then selects out those that, based on significance tests, are correlated with the target. We used the ‘efficient parameters’ setting in tsfresh, whereby features with a high computational cost were excluded. As many of these extracted features will inevitably be highly correlated we subsequently performed PCA to bring the dimensions back down to 32. These 32 PCs were then used as input to GBMs. GBMs are a decision tree ensemble method like random forests. Random forests rely on simple averaging of models in the ensemble, whereas GBMs use boosting, which adds models to the ensemble sequentially (17). At each iteration a new weak base-learner model is trained to focus upon the aspects in the data that have not been well accounted for by the ensemble up to that point. To determine the hyper parameters of the GBMs we drew 200 samples for each fold from the main GBM hyper-parameters. We set a maximum of 300 base learners in the GBM ensemble, but used early stopping with a patience of 10 for selecting the final model.

For the CNNs we made test predictions on the held-out set of cells at each time point between *t* =1 and *t* = 20. For the LSTM and tsfresh applications we instigated 4 checkpoints at *t*=5, 10, 15 and 20.

TensorFlow (version 1.12.0) via Keras (version 2.2.4) was used for the neural network parts of the modeling detailed above and xgboost (version 1.18.1) was used for the GBMs. The tsfresh version used was 0.14.1. Python (version 3.5.2) scripts and jupyter notebooks for the various models are available on GitHub (https://github.com/pharmbio/phil_LNP_modelling).

## Results

The objective of this study was to see the evolution of the classification and regression predictive performance over time for the CNNs applied to the individual time points at test time and to evaluate if performance could be improved by using time-series models that account for time in a more systematic fashion. We assessed this primarily by using the weighted average F1 score for the test set when in classification mode (the higher the better) and using the root mean squared error (RMSE) when in regression mode (the lower the better).

In terms of cell motility our cells moved on average by 9 pixels (1.5 *μ*m) with a standard deviation of 5 pixels (0.8 *μ*m; see Figure 5 for the distribution of movements) between consecutive time points, so there was significant cellular overlap from frame to frame, which boded well for our cell tracking algorithm. However, there were some failures in our tracking algorithm (see discussion).

**Figure 5:**
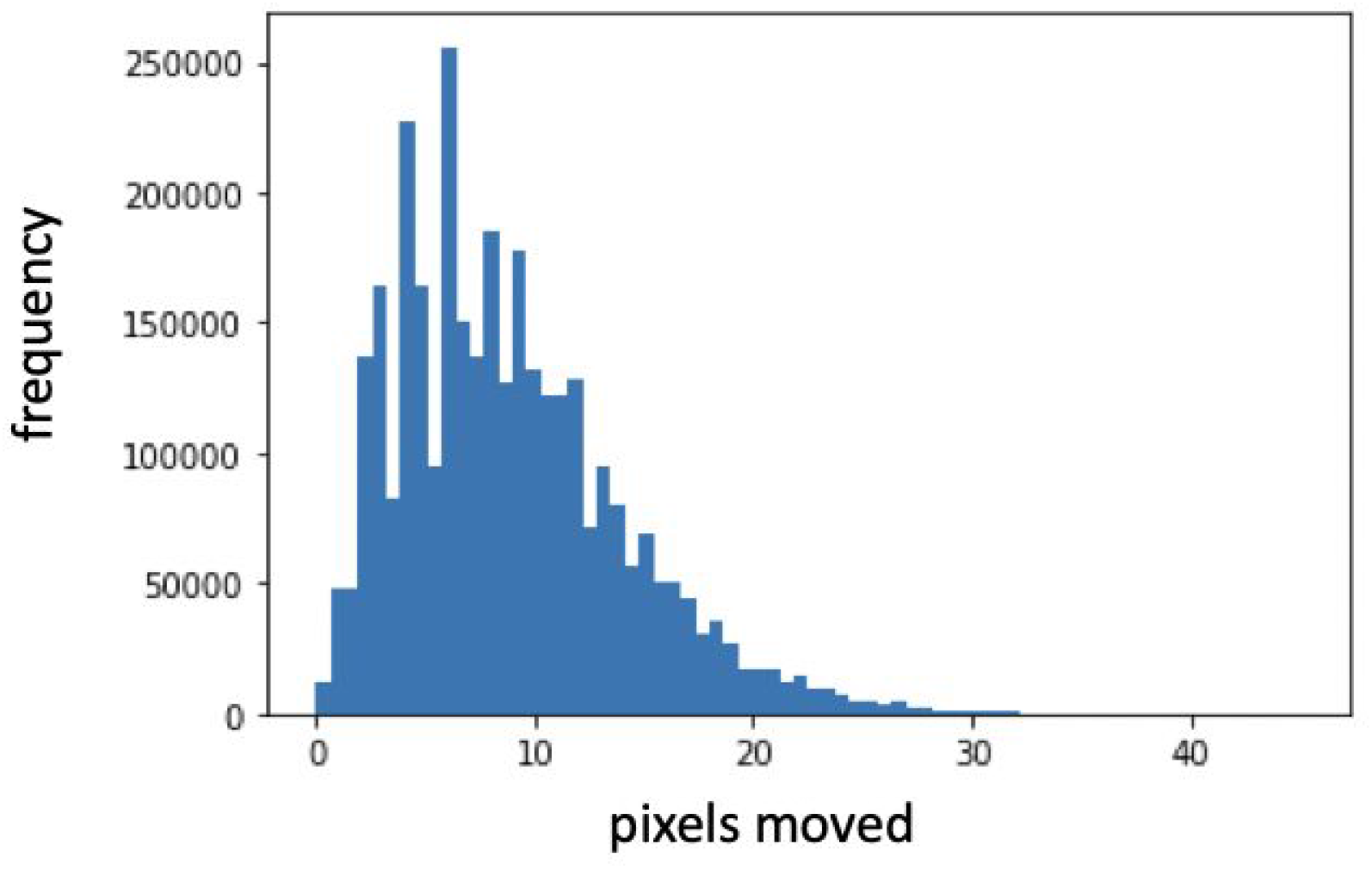
Distribution for the number of pixels moved between each time step for the cells in the training data (mean = 8.92 pixels, 1.49 *μ*m; standard deviation = 5.14 pixels, 0.83 *μ*m).

Based on the CNN explorations with one of the cross validation folds, training for a maximum of 50 epochs with the base frozen and a learning rate of 0.001 was sufficient for both prediction modes (additional adjustments of the base model’s weights, i.e. training with the VGG16 base model unfrozen, or then just it’s final block of convolution operations, did not result in improved performance). For the LSTMs a learning rate of 0.00001 and 30 epochs was selected. The LSTM architectures selected via the grid searches were varying for each fold and each prediction mode, with no clear preferences.

In Figure 6a and 6b we compare the classification and regression performance respectively of applying the CNNs and the time series models to the test data that was withheld during model training. The predictions for each cell were based on the five cross-validation folds. For the time-series models the best model for each fold, from the LSTM or GBM grid searches, was used for making predictions. In classification mode the F1 score increased through time for the CNNs applied to each time point (but plateaued around the tenth time point) and the RMSE decreased continuously through time when in regression mode.

In terms of classification and regression performance, over and above those achieved by the CNNs applied to the data at each time point separately, both the LSTM and tsfresh applications gave significant (randomization test p-values < 0.05) improvements by the twentieth time point and also during some of the previous temporal checkpoints (Table 1). In all temporal checkpoint comparisons these differences were in the direction of improvement (Figure 6), even for the non-significant cases. There were, however, no significant differences between the predictive performance of the LSTM and tsfresh models (Table 1) in either of the prediction modes for any of the checkpoints.

**Figure 6:**
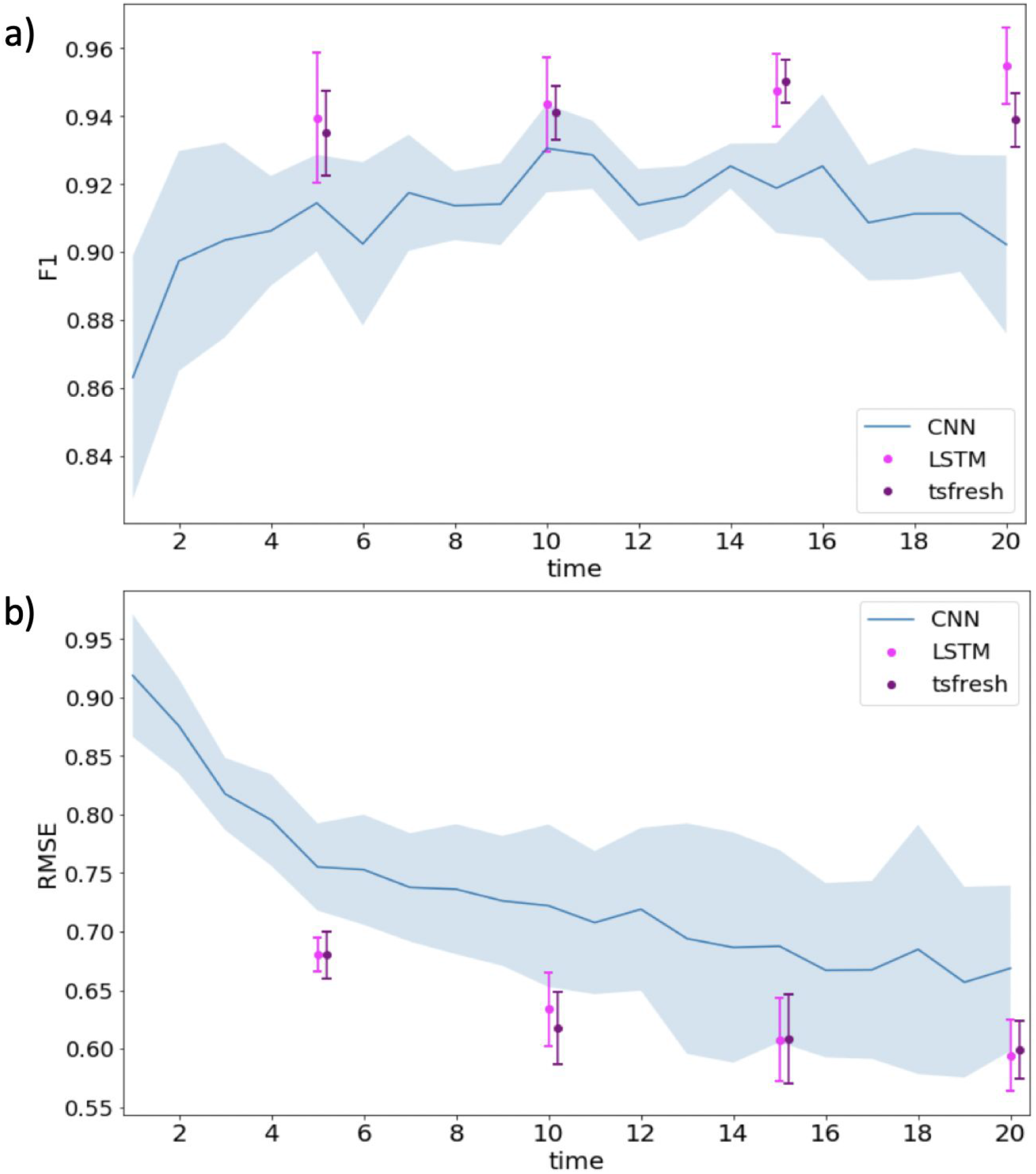
(a) the weighted average F1 score for the held-out test data of cells when in classification mode and (b) the RMSE when in regression mode. The blue line gives the results of the CNN applied to each of the twenty time points and the light shaded blue area shows the uncertainty, one standard deviation above and below the mean (assessed via the five cross-validation folds). The magenta and purple points and lines show the equivalent evaluations at t = 5, 10, 15 and 20 for the LSTM and tsfresh applications, respectively. See Table 1 for an assessment of the significance in the differences between the model comparisons shown here.

**Table 1:** p-values for randomization tests (a cross the five modelling cross validation folds) comparing the prediction results shown graphically in Figure 6. The comparisons are shown in both classification and regression mode, between the CNN alone, the LSTM and the tsfresh applications, for the four temporal checkpoints (t = 5, 10, 15 and 20). The p-values that were significant at the 5%-level are highlighted in red.

| Classification mode |  |  |  |  |
| --- | --- | --- | --- | --- |
|  | t = 5 | t = 10 | t = 15 | t = 20 |
| **CNN vs LSTM** | 0.072 | 0.211 | 0.023 | 0.015 |
| **CNN vs tsfresh** | 0.055 | 0.239 | 0.006 | 0.030 |
| **LSTM vs tsfresh** | 0.718 | 0.734 | 0.665 | 0.077 |
| Regression mode |  |  |  |  |
|  | t = 5 | t = 10 | t = 15 | t = 20 |
| **CNN vs LSTM** | 0.007 | 0.022 | 0.086 | 0.039 |
| **CNN vs tsfresh** | 0.015 | 0.008 | 0.096 | 0.017 |
| **LSTM vs tsfresh** | 0.965 | 0.512 | 0.971 | 0.857 |

For the LSTM and tsfresh applications we show in Figure 7 a plot of the true versus predicted GFP values for the test cells (sorted by their GFP expression level) when in regression mode at the *t* = 20 temporal checkpoint.

**Figure 7:**
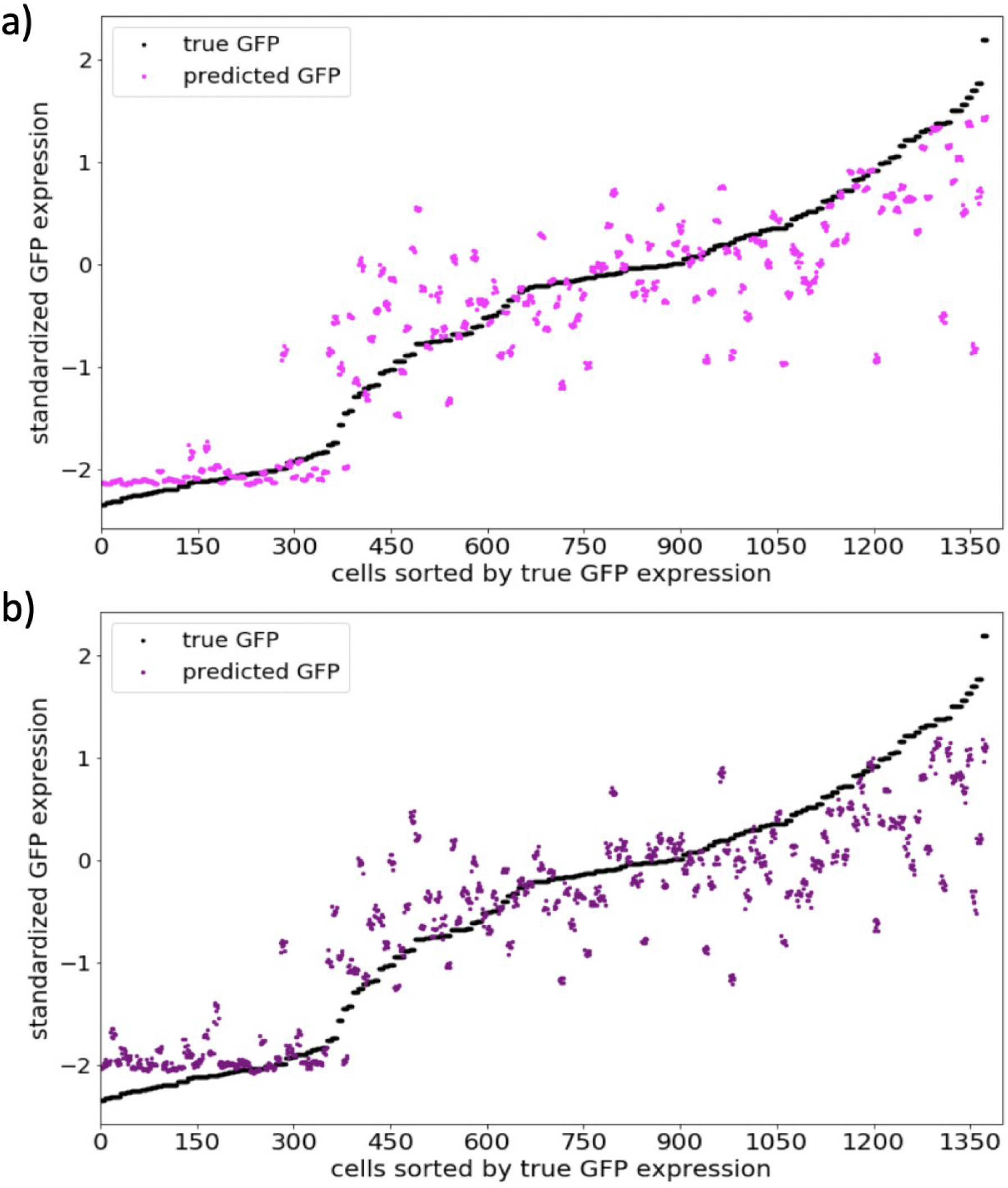
(a) regression results for the LSTM from Figure 6b at the t = 20 temporal checkpoint, where the black dots show the true GFP value for the test cells (sorted by their GFP value) and the magenta dots their predictions. (b) the equivalent results to (a) for the tsfresh-based application with the predictions shown by purple dots. There were 172 cells in the tests set, but as we applied 8x test-time augmentation, there were essentially 1376 test objects. The predictions were based on the averages across the five cross-validation folds.

## Discussion

In this study we have shown that it is possible to predict GFP expression in advance from cell-level microscopy image data by capturing information about the morphologies of the cells and their interactions with the LNPs. This was also possible to do with a relatively small dataset. Furthermore, we have also shown that explicitly accounting for the temporal dynamics, using either LSTMs or tsfresh, led to significantly improved predictive performance. Although neither of these two time-series modelling approaches shined out as being superior for our predictive purposes.

The information richness in single cell biology presents both a major challenge and an opportunity (18). The richness comes from the dearth of information and the challenge comes in terms of how to process and make sense of this volume of information and to organize it into a coherent whole. For the latter, if we are able to identify the most relevant data points early in an experiment, more efficient experiments can potentially be designed, less data is acquired and computational resources can be used more efficiently. Biologically this also enables a different sort of analysis that has not previously been possible when using end-point assays.

Despite the fact that our CNNs were not explicitly provided with temporal information, the increase in F1 score (for classification) and reduction in RMSE (for regression) through time suggest that our CNNs might be able to organize the information temporally, such that the cell features from later time points were more informative for the predictive tasks at hand than the earlier ones. This behavior has previously been shown by Eulenberg et al. (18) where CNNs were used to reconstruct cell cycles and disease progression through time.

When training CNNs with transfer learning, unfreezing the entire base model, or only the later convolutional layers, often yields improved predictive performance (4, 19). The fact that we did not find this to be the case here may be due to the fact that we only took the first three (of five) blocks of convolution and pooling from the VGG16 network. The earliest layers of CNNs extract information common to all images, such as edges and blobs, whereas later layers become more specific to the task at hand. Unfreezing our base model, and fine tuning the weights in these early layers, is thus less necessary.

For the regression models, although the increasing trend in GFP expression was followed for the predicted values, there was a large amount of variance and some clear outliers (especially for some of the cells with a high GFP expression level, Figure 7). These outliers were common to both the LSTM and tsfresh approaches. To rule out that this was not caused by failure in the cell tracking, we visually inspected the 172 time lapse videos for the test cells, and confirmed that this was not the source of the problem. Of the 172 videos, however, we found 14 failures (8%), i.e. cases where the focal cell was lost. Of these, 6 (3%) did not have the focal cell within the cropped window for creating the data used in the models.

It is often difficult to determine what CNNs actually learn and many people (including scientists) still view deep learning as something of a ‘black box’ methodology. A great amount of scientific effort, however, has recently been put into ‘explainable AI’ (20) which aims to both interpret and visualize the details that such models have focused upon in order to attain their [often impressive] predictive performance. Such interpretations and visualizations for LNP applications would be a good subject for future studies. Discriminating what are the important features in the input image or time series can lead to important biological findings. Nevertheless it may still be somewhat (or even much) easier to trace back through trained models that are based upon more directly interpretable features, such as those using time series feature extraction (e.g. from tsfresh) atop more conventional spatial feature extraction (e.g. those computed by the well known CellProfiler (21) software package). As we did not find significant differences between the LSTM and tsfresh based approaches this consideration would argue for the continued use of tsfresh. However an argument in favor of the LSTM based methods is the fact that it is also possible to train the CNN-LSTM models in an end-to-end fashion (i.e. whereby the weights in both the CNN feature extractor and the LSTM can be further fine-tuned in unison to potentially improve predictive performance). Such fusing is not possible for the CNN-tsfresh methods. In cell biology jointly trained CNN-LSTM networks have previously been used, for example, in the detection of mitotic events from phase contrast microscopy images (27).

Other methods, besides those we chose to focus upon in the current study, have also proven insightful for elucidating cell morphological dynamics from time-lapse microscopy, such as the combination of temporally constrained combinatorial clustering, gaussian mixture models and hidden markov models used by Zhong et al. (22). Another interesting time-series approach, not specific to cell dynamics, proposed by Wahlström et al. (23), uses autoencoders to reconstruct future (as opposed to the current) images combined with an embedded dynamic model. Defined in this way, the goal is to minimize a dual cost function (with one component for the future image reconstruction and the other for the prediction error). Their model essentially learns how to compress the image data into components relevant for the underlying temporal dynamics. Their method, however, requires a realistic model for the dynamics of the system under study. The methods we focussed upon in our study are more flexible in the sense that knowledge of how such an underlying model should be composed is not required.

Despite our relatively small dataset size we still managed to achieve good predictive performance and showed that it was possible, using transfer learning, to train models on as few as 774 cells. This is encouraging for studies when it is not possible to obtain a large number of cells. However, with the limited number of cells in our study, it was necessary to perform cross-validation and testing at the cell-level after the cells had been randomly shuffled. A larger dataset would make it possible to cross-validate and also draw conclusions at the well level. Another opportunity for future work would be to include a nuclear counterstain (imaged only in the first time frame; to avoid detrimental effects upon the cells) as a basis for seeding the watershed algorithm in the first time point. This would avoid the need for manual centering at this time point - a procedure that was feasible for our current dataset, where we had less than one thousand cells, but would not be so for larger datasets. Additional, sporadic, nuclear counterstain images through time could also help improve our tracking algorithm, as would accounting for previous cell shape and motility behaviour. Such motility behaviour could also be incorporated into our models. Kimmel et al. (24) have previously used a combination of LSTMs and 1D CNNs for cell motility discrimination and prediction (based on cell track data). Although our CNNs and time-series modeling approach may partially learn this motility behaviour indirectly (e.g. through dynamic changes in the geometries of the cells), including this information directly may improve predictive performance. Another statistic of interest to include in future models is the density of cells in the vicinity of the focal cell. Again, this is something that our models may have learnt indirectly, where, for densely packed areas, our data preprocessing crops would have partially included the closeby cellular neighbours.

## Code availability

Code scripts and jupyter notebooks, written in python, for the modeling performed in this paper are available on GitHub (https://github.com/pharmbio/phil_LNP_modelling).

## Acknowledgements

This project was financially supported by the Swedish Foundation for Strategic Research (grant BD15-0008SB16-0046) and the European Research Council (grant ERC-2015-CoG 683810). Thanks go also to Marco Capuccini, Alex Danis, Anindya Gupta, Ankit Gupta, Anders Larsson, Thomas Schön, Oliver Stein, Robin Strand and Niklas Wahlström for their help and advice.

